# Single-Cell Multi-Modal GAN (scMMGAN) reveals spatial patterns in single-cell data from triple negative breast cancer

**DOI:** 10.1101/2022.07.04.498732

**Authors:** Matthew Amodio, Scott E Youlten, Aarthi Venkat, Beatriz P San Juan, Christine Chaffer, Smita Krishnaswamy

**Affiliations:** Yale University Department of Computer Science; Garvan Institute of Medical Research, Darlinghurst, NSW, Australia; Yale University Computational Biology and Bioinformatics; St Vincent’s Clinical School, UNSW Medicine, UNSW Sydney, NSW, Australia; The Kinghorn Cancer Centre, Darlinghurst, NSW, Australia; Yale University Department of Genetics

## Abstract

Exciting advances in technologies to measure biological systems are currently at the forefront of research. The ability to gather data along an increasing number of omic dimensions has created a need for tools to analyze all of this information together, rather than siloing each technology into separate analysis pipelines. To advance this goal, we introduce a framework called the Single-Cell Multi-Modal GAN (scMMGAN) that integrates data from multiple modalities into a unified representation in the ambient data space for downstream analysis using a combination of adversarial learning and data geometry techniques. The framework’s key improvement is an additional diffusion geometry loss with a new kernel that constrains the otherwise over-parameterized GAN network. We demonstrate scMMGAN’s ability to produce more meaningful alignments than alternative methods on a wide variety of data modalities, and that its output can be used to draw conclusions from real-world biological experimental data. We highlight data from an experiment studying the development of triple negative breast cancer, where we show how scMMGAN can be used to identify novel gene associations and we demonstrate that cell clusters identified only on the scRNAseq data occur in localized spatial patterns that reveal insights on the spatial transcriptomic images.

## 1 Introduction

Integrating data gathered from different sources is a critical challenge in computational genomics. Currently there are several single-cell technologies including RNA-sequencing, ATAC-sequencing, Hi-C, ChIP-seq, and CITE-seq as well as proteomic technologies such as CYTOF, imaging CyTOF, and MIBI that offer complementary cellular information [1, 2, 3, 4, 5, 6, 7]. Of these modalities only a fraction are available as simultaneous measurements—often with quality degradation factors such as reduced gene dimensions, lower throughput, and increased noise. The remaining measurements must be done on distinct cellular subsamples from the same population. This is the key problem that we tackle in this manuscript: the prediction of missing or non-simultaneous modalities in order to generate a more complete set of features. Thus given modalities like scRNA-seq, scATAC-seq, and spatial transcriptomics measured separately on different cells (from the same population), scMMGAN generates a complete set of simultaneous measurements for downstream analyses.

Aligning the separately-measured data computationally has many advantages over analyzing the data modalities individually. Combining data modalities allows us to leverage the advantages of each and mitigate the disadvantages. For example, combining a modality with a higher signal-to-noise ratio like proteomic CyTOF measurements with one that has a lower signal-to-noise ratio like single-cell RNA sequencing gives us the opportunity to resolve cell populations to a finer degree in the noisier dataset [7]. Even more compellingly, combining modalities allows us to measure variables only available in one domain combined with variables only available in another domain, thus simulating jointly measured technologies.

We base our method on the framework of cycle-consistent generative adversarial networks (CycleGANs) [8, 9, 10, 11]. In GAN-based domain adaptation frameworks, a generator network is trained to map data points of one modality into data points from another modality. During training, a discriminator is used to ensure that generated points are sampled from the high dimensional distribution representing the second modality. In CycleGANs, there are two back-to-back generators, one going from the first modality to the second modality, and another going from the second modality to the first. A reconstruction error enforces that the result of two back to back domain adaptations results in the original distribution again i.e. that the generators are inverses of each other over the regions of the data spaces where there are training points.

While CycleGAN frameworks can successfully generate points in each modality, the *mapping* they produced is not constrained enough. For instance in the original CycleGAN paper, images of zebras were mapped to images of horses, but nothing ensured that the background would be unaltered. While this may not be detrimental for natural image applications, it can be untenable for scientific applications where the scRNAseq measurement must *correspond with and corroborate* the scATAC-seq measurement. Noting this key weakness, in earlier work, we proposed the use of a *correspondence loss* and gave anecdotal examples on flow cytometry panels with overlapping measured markers [12]. However, here we both extend the application to multi-modal integration and specify a more powerful, generally applicable correspondence loss: the *geometry preserving loss.* This loss enforces that the diffusion geometry, performed with a new kernel designed to pass gradients better than the Gaussian kernel, is preserved throughout the mapping. We note that this loss can be utilized even in cases where no measurements overlap.

We demonstrate the power of aligning data modalities with scMMGAN on a wide array of data types. We start by validating it on datasets where simultaneous measurements are available and use those as ground truth in evaluations. Then we use scMMGAN to perform a thorough investigation into a novel triple negative breast cancer dataset, where we have cells from the breast cancer culture HCC38 xenografted into mice and allowed to metastasize from the primary to secondary tumor locations. We show that scMMGAN can infer spatial locations of cellular structures.

## 2 The scMMGAN Model

### Architecture

The scMMGAN framework is depicted in Figure 1a. Each pair of data domains or modalities has a pair of generator networks that map in either direction between them, forming a many way multi-modal mapping. For a generator mapping from Domain *i* to Domain *j* which we denote *G_ij_*, it functions as a traditional GAN guided by a discriminator in Domain *j* which we denote *D_j_*. The discriminator tries to distinguish between samples from the real data for that domain *x_j_* and samples from the generator *G_ij_* (*x_i_*), while the generator tries to fool the discriminator. They alternate trying to optimize the following minimax objective:

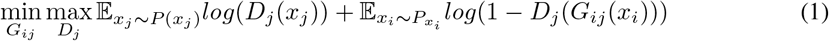

**Figure 1:**
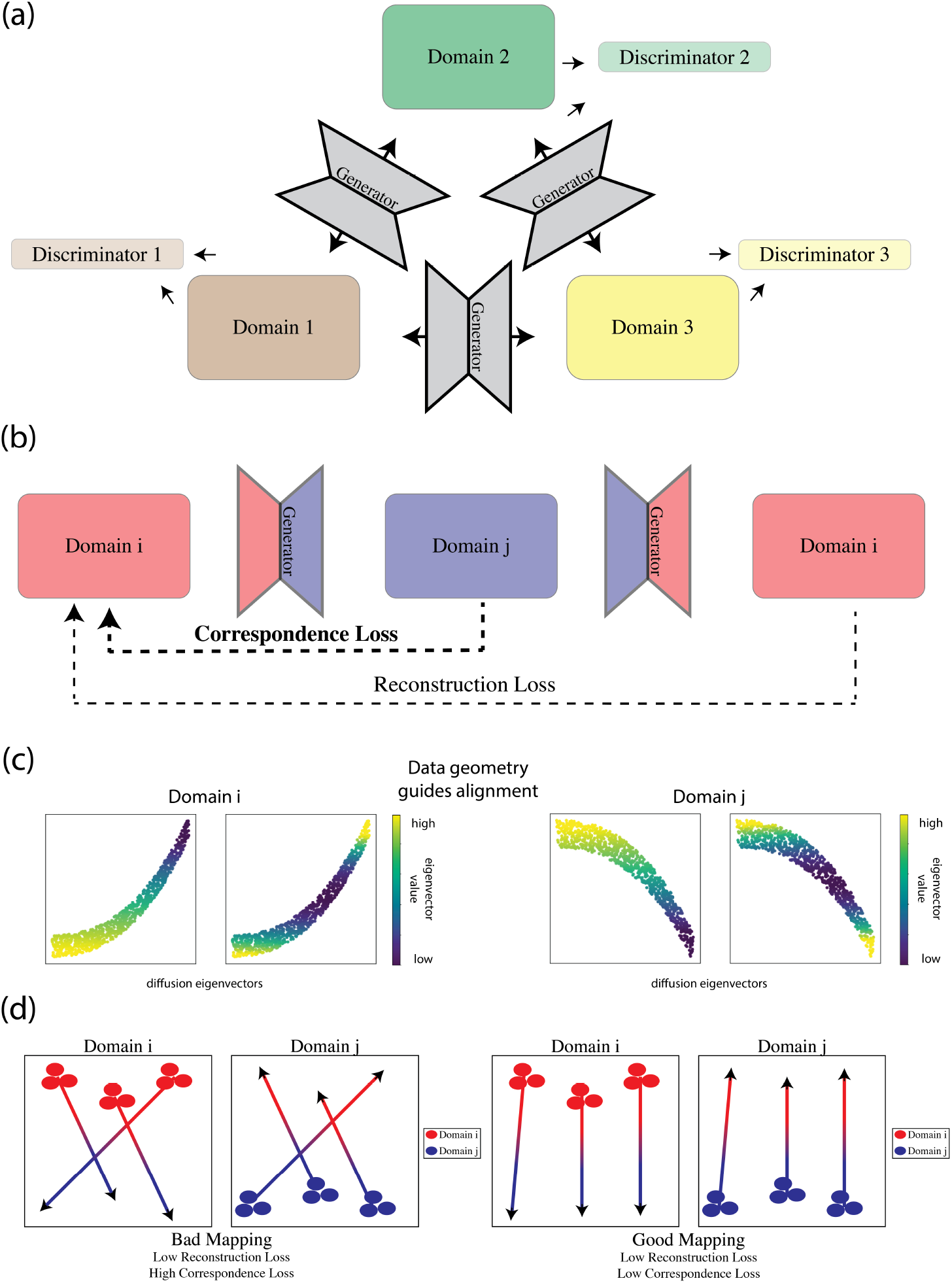
(a) The scMMGAN architecture mapping between multiple domains, each consisting of a pair of generators and discriminator. (b) In addition to the discriminator loss, there are two additional losses within each domain. (c) Hypothetical demonstration of the data geometry guiding alignment through the correspondence loss. In the depicted space, data in the two domains have been shifted and rotated, but the intrinsic data geometry as is preserved with the values of the diffusion eigenvectors. (d) Hypothetical illustration of a bad mapping that is invertible (has low reconstruction loss) but doesn’t align analogous representations (has high correspondence loss) and a good mapping that is both invertible and aligns analogous representations. In the situation where minimally changing the value of genes is preferred, the mapping on the left unnecessarily changes the value of the gene on the x-axis.

In addition to this loss of the discriminator guiding the generator to transform its input modality into the output modality *L_GAN_*, there are two other terms in the loss that ensure the learned mapping is informative and meaningful. These are depicted in Figure 1b. The reconstruction loss *L_r_* is the mean-squared-error (MSE) between the original data *x_i_* and the composition of the two paired generators between the domains *i* and *j*:

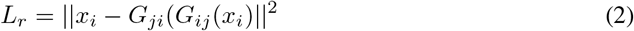

The correspondence loss *L_c_* imposes a constraint on a single point’s representation in Domain *i* and Domain *j*, as opposed to the reconstruction loss which imposes a constraint on points within the same domain:

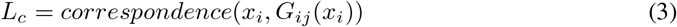

The motivation for the correspondence loss comes from the fact that previous models using cycle-consistency for domain mapping with GANs included only two restrictions: (1) that the generators be able to reconstruct a point after it moves to the other domain and back, and (2) that the discriminators not be able to distinguish batches of true and mapped points. The generators can accomplish these goals in many different ways, including by learning arbitrarily complex mappings: as long as they align the two data manifolds at a distribution level. The family of paired inverse functions that can match the target distributions is large, and with existing frameworks, the particular pair that results from training is determined by the vagaries of random weight initialization and mysterious biases in gradient descent.

The GAN loss is a probability-distribution matching objective [13]. In previous work it has been proven that under certain optimality conditions that a GAN discriminator provides a Jensen-Shannon divergence between the true and generated distributions [14]. A Wasserstein GAN (WGAN), on the other hand contains modifications that result in a wasserstein distance being provided by the discriminator [15].

However simply matching probability distributions can result in incoherent cell states (Figure 1d). A key insight we bring is that distributions must only be matched within correspondence constraints. These correspondences are essentially invariances in the underlying system that are reflected in every modality. In our previous work we used customized correspondence losses for each dataset. However, here we note that when matching single cell data we can use a nearly universal constraint—that of manifold geometry preservation.

While our model incorporates signal from a data geometry loss into a larger framework, the data geometry is too rigid to be used on its own to guide alignment. It is heavily influenced by the properties of the domain data space, and thus when the two domains are very different it does not allow for sufficient flexibility in changing the shape of the distribution. Methods that use only the data geometry struggle to align domains that are significantly different [16]. An ideal mapping would have both the flexibility of a mapping that matches the probability distribution (like the GAN does) but preserves the data geometry as much as possible while doing so. By combining the existing GAN-based loss and a data geometry loss, the network can balance the tradeoff between these goals.

We thus introduce a correspondence loss that ensures the mappings have pointwise as well as distributional alignment by preserving the data geometry through the learned mapping. To do this, we use the diffusion map representation of the original data.

Here we give a brief overview of diffusion maps. Diffusion maps are a kernel-based method frequently used in manifold learning to produce low-dimensional embeddings that preserve intrinsic structure in the data [17, 18]. The eigenvectors of the diffusion operator form an embedding where Euclidean distances correspond to diffusion distance, or the probability of getting from one point to another via random walk, on the original manifold [19]. Because these new coordinates represented by the diffusion eigenvectors abstract away much of the data-domain specific properties, they present a way of ensuring the underlying data geometry is preserved in the mapping.

By calculating the eigenvectors of the diffusion operator for the points in their original domain *ϕ*_*x*_1__ and the eigenvectors of the diffusion operator for the points after being mapped to the other domain *ϕ*_*G*(*x*_1_)_||_2_, we can directly compare the *i^th^* eigenvectors to enforce that intrinsic data geometry, as measured by diffusion, be preserved by the mapping. For further detail about the calculation of *ϕ*, please see the experimental procedures section later.

The correspondence geometry loss then penalizes the L2-distance between the two representations of each point:

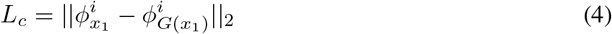

By enforcing this loss in the scMMGAN framework, we ensure that the intrinsic structure of the data is preserved in the otherwise under-constrained GAN setting.

## 3 Experimental Results

### 3.1 Mapping between spatial, scRNA-seq and proteomic data

As an initial validation of scMMGAN, we utilize simultaneously measured multimodal data from the newly developed DBIT-seq (deterministic barcoding in tissue) technology as ground truth. DBIT-seq uses deterministic barcoding in tissue for spatially resolved measurements of both transcriptomics and proteomics [20]. Thus, in DBIT-seq, three things are measured jointly on every cell: a scRNAseq profile, a protein profile, and spatial coordinates. The system being studied in this data is that of mouse embryos, particularly focused on early tissue development and organogenesis.

Often, in transcriptomic/proteomic alignment problems, no “ground truth” is available because each technology measures a distribution of cells in a destructive process. As a result, models like scMMGAN that learn to map between two distributions without needing point-wise pairings are called unsupervised alignment models. We design an experiment with this data to show how scMMGAN could have been used to obtain this information without needing them to be measured jointly. We treat the spatially located scRNAseq data and the spatially located protein data as two separate measurements and learn to map between them. We then utilize the fact that they were measured jointly and that some of the columns in each dataset are related (corresponding genes and proteins) to evaluate the accuracy of the learned mapping. We compare against both an autoencoder-based alignment method (CMAE) and a standard CycleGAN without a correspondence loss [9, 21]. For detailed descriptions of the model architectures, please see the section on experimental methods.

Figure 2 shows example results of scMMGAN and baseline models on this data. Plotted on the given spatial coordinates, we show the ground truth transcriptomic value along with generated proteomic values. There we see scMMGAN best models the ground truth data. We further evaluate scMMGAN’s performance on this application quantitatively. To quantify the aspect of the generated distribution matching the target distribution as a whole, we employ the metric Maximum Mean Discrepancy (MMD), a distance defined on distributions frequently used in both deep learning and biological contexts to distinguish between distributions [22, 23, 24]. To quantify the aspect of preserving information about the individual observation through the alignment, we use correlation between columns in the transcriptomic space and the proteomic space known to correspond to the same gene. Since these values correspond to the same gene, we would expect there to be a correlation between a point’s value before mapping and its value after mapping.

**Figure 2:**
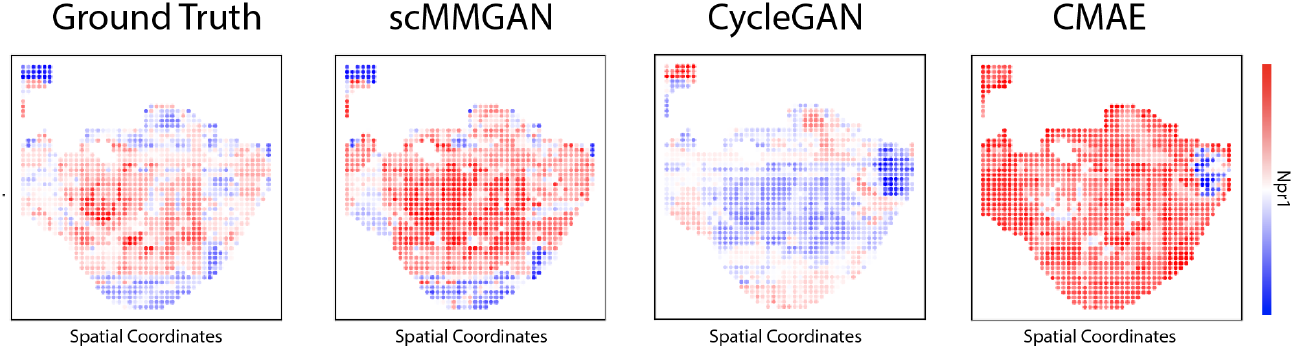
On the DBIT-seq data, shown are corresponding proteomic and transcriptomic expression for the gene shown. The x-axis and y-axis plotted are the measured spatial coordinates taken directly from the data. The ground truth transcriptomic values are plotted alongside the generated proteomic values for each model, where we see scMMGAN best model the data.

These scores confirm quantitatively what we saw graphically in these experiments (Table 1). All models are able to accurately match the target distribution (low MMDs), with very similar performance consisting of each model’s one or two standard deviation interval overlapping. However, when looking at the preserved correlation, we see scMMGAN achieved the best alignment with an average correlation of *r* = 0.154 between columns known to correspond. We note that the absolute value of this correlation is relatively low compared to other datasets, and this is due to limited amount of shared correlation in the underlying “ground truth” pairings of points that are jointly measured.

**Table 1:** Evaluation of each model on the DBIT-seq data. While the MMDs are close for each model (meaning the predicted distribution resembles the ground truth distribution), scMMGAN is significantly more accurate at preserving correlation between columns known to correspond. The best values are bolded.

| DBIT-seq | scMMGAN | CycleGAN | CMAE |
| --- | --- | --- | --- |
| $\text{MMD}(G(x_1), x_2)$ | **0.072** +/- 0.001 | 0.078 +/- 0.006 | 0.082 +/- 0.001 |
| $\text{MMD}(x_1, G(x_2))$ | **0.060** +/- 0.001 | 0.066 +/- 0.002 | 0.079 +/- 0.001 |
| $\text{Correlation}(G(x_1), x_2)$ | **0.155** +/- 0.006 | -0.026 +/- 0.021 | 0.003 +/- 0.082 |
| $\text{Correlation}(x_1, G(x_2))$ | **0.152** +/- 0.014 | -0.088 +/- 0.074 | -0.012 +/- 0.066 |

#### Unique vs Common Information in Measurement Modalities

The scMMGAN is a generative framework, but when used in non-standard ways it can become a tool for analysis in addition to faithful generation. When measuring two aspects of a biological system with two different technologies, some of the information might be shared between the two modalities while other information might be unique to one of the modalities. For example, when mapping between a modality that measures the whole transcriptomic space like scRNAseq and one that measures only a subset of the proteomic space, we would expect for the genes with corresponding proteins to be more easily modeled than the genes without them.

We design our experiment as follows to test this on the transcriptomic and proteomic measurements in the DBIT-seq data (summarized in Figure 3, with further mathematical detail in the methods section later). We train the model augmented with random noise input and then evaluate it on mapping the same points in proteomic space to the transcriptomic space, except with different random noise samples. We then calculate the variance of the different predicted values for each transcript count and for each given point in proteomic space. Then, the mean across all points gives us a measure of the uncertainty associated with a given transcript measurement. To factor out the influence the magnitude of counts of a given transcript would have on variance, we scale by the variance in the raw dataset for each one. We also filter out lowly expressed genes. We can then compare the stochasticity as measured in this experiment of the genes which have a corresponding proteomic measurement and those that do not.

**Figure 3:**
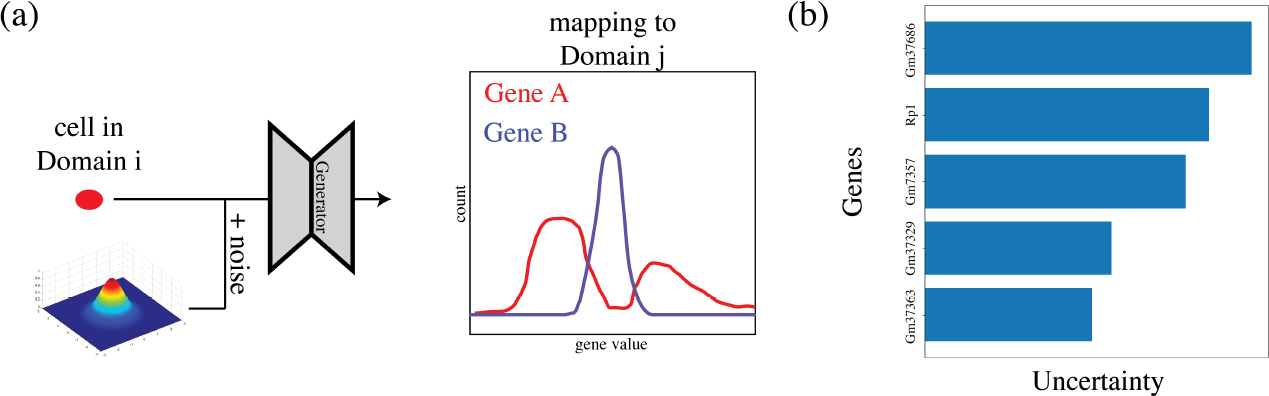
(a) A depiction of how scMMGAN can be used to quantify how much uncertainty is associated with the mapping to each gene. A particular cell is mapped from Domain *i* to Domain *j* along with various different noise samples. The mapped values of Gene A change significantly with the noise, while the mapped values of Gene B change little for this cell. We interpret this as a quantification of how much information there is about each gene in Domain *i*. (b) The genes identified by scMMGAN to have the most uncertainty associated with the mapping, and thus have the least common information with the proteomic measurements in this dataset.

Just as expected, we find that the average variance of genes with an analogous proteomic measurement is 0.026 while the average variance of genes without an analogous proteomic measurment is 1.419. This is a logical result as the relationship between transcript counts with corresponding proteomic measurements is more straightforward to model and can thus be done with more certainty. This corroborates our understanding of the process at work in this multimodal setting.

Furthermore, this analytic process with scMMGAN allows us to inspect which genes are modeled with the most and least uncertainty and thus provide the most unique information and most common information with respect to the other modality. Unsurprisingly, the genes with the least uncertainty are related to early embryo development in mice, as that is the system being studied: for example the three least are Gm5049, Gm37500, and Gm33051. Meanwhile, the genes with the most uncertainty are GM37686 and Rp1. It is possible that genes with higher observed uncertainty could also be of interest to the research, for example Rp1 which is involved in the development of the retina and the study was focused on early tissue development [25]. The information that these genes had high uncertainty can help guide future experimental design decisions that would lead to the selection of proteins to measure that would allow for better alignment of these measurements.

In this way, scMMGAN can help provide insights into the system being studied as well as into experimental design and decisions.

### 3.2 Mapping between scRNAseq and ATAC-seq data

We next perform an experiment on data consisting of paired ATACseq and RNAseq measurements on the same cells. As with the previous experiment, since these two technologies both measure values related to a particular gene (chromatin availability for ATACseq and transcript expression for RNAseq), we can expect there to be some correlation between the two spaces in their values for that gene, as in the prior case. The dataset we use comes from a public human blood dataset of granulocytes removed through cell sorting from PBMC of a healthy donor [26].

A qualitative assessment of the results via plots of the output are shown in Figure 4, with the ground truth ATAC value plotted in the first column, and then the generated corresponding RNAseq value in the subsequent columns. As before, scMMGAN’s output best matches the ground truth. For the other models, while they have the right amount of activation for each gene at a distribution level, they are inaccurate in terms of alignment at a point level (some populations have been inverted).

**Figure 4:**
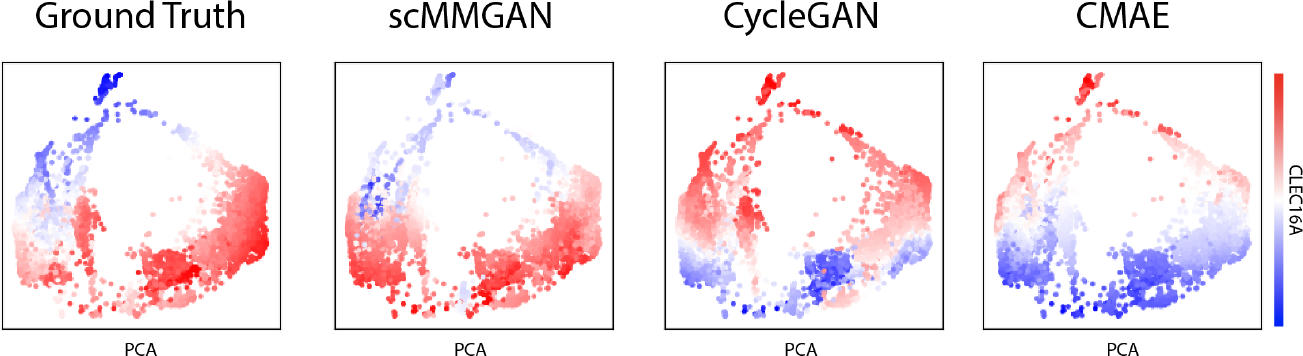
Ground truth values for held-out cells and the predictions for each model on the experiment mapping between ATAC and RNA sequencing. scMMGAN’s output matches the ground truth most accurately compared to the other models, which inverted populations through the mapping. Coordinates shown are from the first two PCA dimensions.

Confirming this quantitatively, as seen in Table 2, while all of the models perform adequately at matching the ground truth at a distribution level (as seen by their low MMDs), a significant difference can be seen when evaluating them at a point-wise level. scMMGAN’s predictions have an average correlation with the ground truth of *r* = 0.336 while the others are all essentially uncorrelated (0 is near the middle of each models’ 1-σ interval).

**Table 2:**
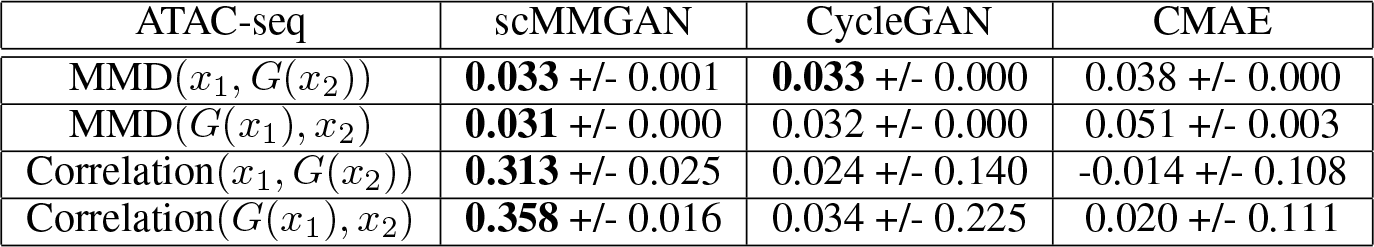
Evaluation of each model on the ATAC/RNA seq data. The MMDs for each model are close as each models the ground truth at a whole-distribution level. scMMGAN is the only model whose predictions preserve the known correlation though, because its alignment is also accurate point-wise. The best or tied-for-best values are bolded.

### 3.3 Integration of Triple Negative Breast Cancer Data

Here, we apply scMMGAN to a dataset that comprises of a human xenograft model of triple-negative breast cancer (MDA-MB-231) with transcriptomic measurements jointly in both a spatial RNAseq modality and a scRNAseq modality lacking the spatial information. The study consists of the MDA MB 231 cell line grown in mouse models, with the measurements taken from primary site tumors in the tissue from the mammary gland. While the experimental models are replicates, they are different organisms and thus introduces an additional source of noise for the alignment.

Each of the two measurement modalities produce transcriptomic measurements, but each also has advantages and disadvantages. The spatial RNAseq provides the ability to analyze the physical structure of the tissue sample, and localize behavior to different regions of it via (x, y) spatial coordinates. As a drawback, though, each spatial coordinate is bigger than the size of a single cell, and as a result the transcriptomics are estimates of groups of multiple cells. For example, if a cell of one type that is expressing gene A and a cell of another type that is expressing gene B are spatially adjacent, this technology would observe gene A and gene B being expressed together, even if they are never jointly expressed in a single cell.

In contrast, the scRNAseq provides the usual single-cell granularity of measurements that would be able to distinguish between the expression of each cell. By mapping the spatial data to the scRNAseq space, we are in essence imputing it into single-cell resolution. However, the scRNAseq does not have spatial orientation with respect to the original tissue sample. Thus, to combine the best of each modality (spatial information at the single-cell level), we use scMMGAN to integrate them by mapping a point from the spatial RNAseq domain to the scRNAseq domain and considering its aligned representation of its original spatial coordinates and its generated scRNAseq expression values.

The spatial RNAseq dataset consists of a tissue sampled across 1170 spatial coordinates, each coordinate with measurements on 20092 genes. Four different scRNAseq samples were obtained from cancerous primary site tissue (from different mice), each measured across the same 20092 genes, and consisting of 7606, 5118, 8163, and 7591 cells, respectively.

While the scRNAseq and spatial RNAseq data are both transcriptomic technologies measuring gene profiles, and thus their dimensions have the same meaning, the two datasets cannot be analyzed together as is. In Figure 5a, we see that the two data distributions are completely non-overlapping prior to the use of scMMGAN. Since the raw data is completely separable in the joint space, any downstream analysis would only be able to pick up on the difference between the two modalities and not the differences between cells within them. For an integrated analysis using information from both of them, we need the aligned output from scMMGAN (Figure 5b).

**Figure 5:**
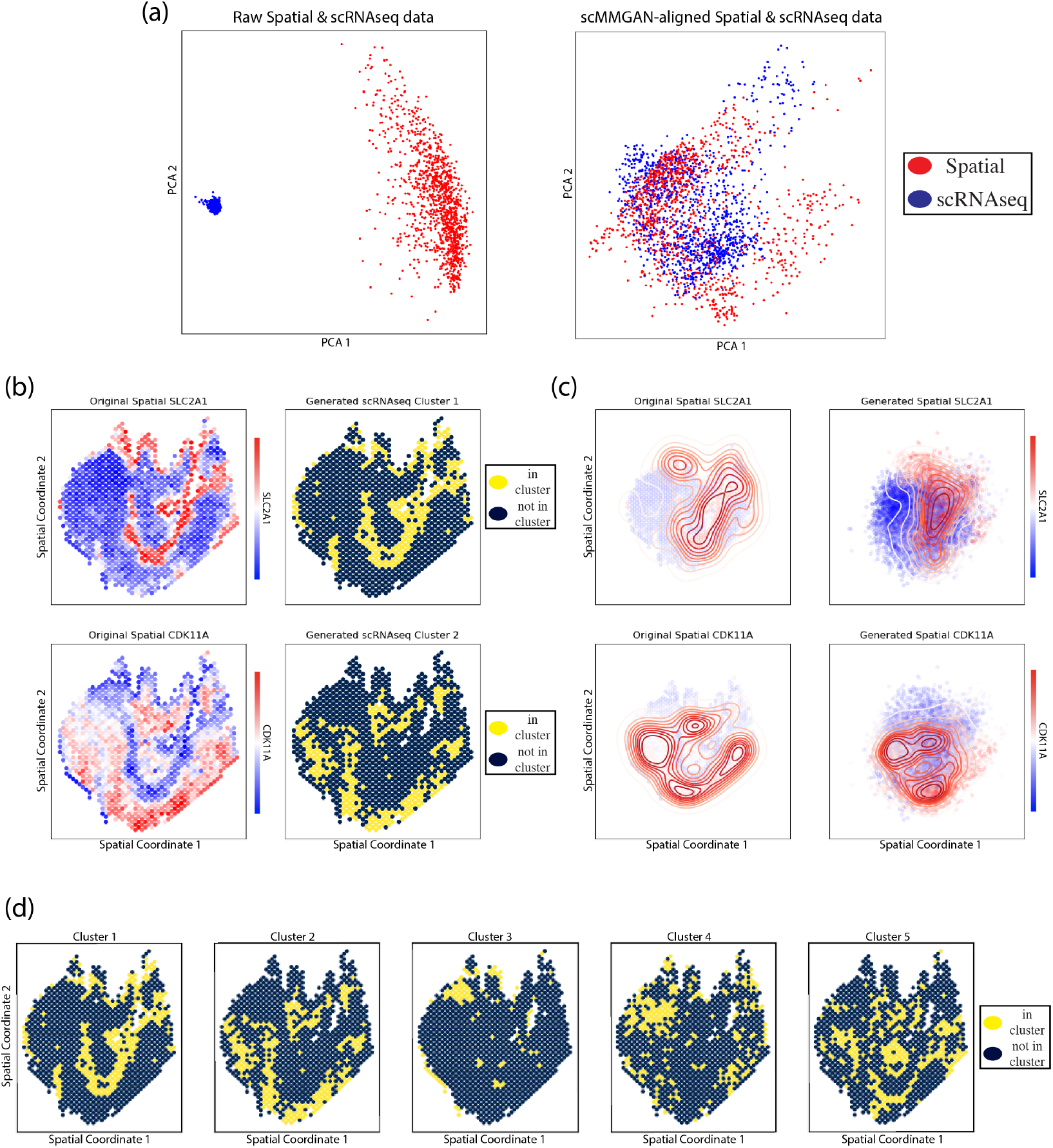
(a) Plotted are the PCA coordinates of the gene expression values from the two distributions. In the raw data, the spatial RNAseq and scRNAseq are not directly comparable, as they are entirely separable. After mapping with scMMGAN, they are aligned and comparable with downstream analysis. (b) Mapping spatial RNAseq to scRNAseq, clustering the generated scRNAseq values, and then plotting the cluster by the measured spatial coordinate on the x-axis an y-axis. (c) Generated spatial RNAseq data from scRNAseq, including generated spatial coordinates. Same coordinates as previous plot. (d) All generated clusters mapped to the spatial RNAseq space. Same coordinates as previous plots.

We analyze the scMMGAN alignment by taking the spatial RNAseq, mapping it to the scRNAseq space, and then clustering the generated scRNAseq data (Figure 5b). We use spectral k-means clustering with a selected parameter of *k* = 5, and we then plot the clusters according to their original spatial coordinates. As we see, these scRNAseq clusters preserve spatial patterns seen in the coordinates, demonstrating our ability to make new spatially-informed conclusions through analyzing the generated scRNAseq data in conjunction with the original spatial coordinates.

In Figure 5c, we look at the opposite mapping direction of taking the scRNAseq data and generating spatial RNAseq with it. By mapping scRNAseq points to these generated coordinates, we can see spatial organization of particular cell types. For example in this figure, we plot the generated spatial coordinates for cells high in SLC2A1 and CDK11A and see spatial differentiation between these two types of cells. We show all of these clusters plotted in Figure 5d.

Now that we see scMMGAN has learned to map data between the two modalities in a way that preserves gene signals, we next compare this to the alternative alignment models. In Figure 6, we show the results of the learned mapping from spatial RNAseq to scRNAseq for each model. We first note that each model was able to generate a distribution that accurately matched the target distribution, an observation we will demonstrate quantitatively later. However, the alternative approaches to scMMGAN achieved this result by aligning a given spatial RNAseq gene profiles to a scRNAseq observation that is very different.

**Figure 6:**
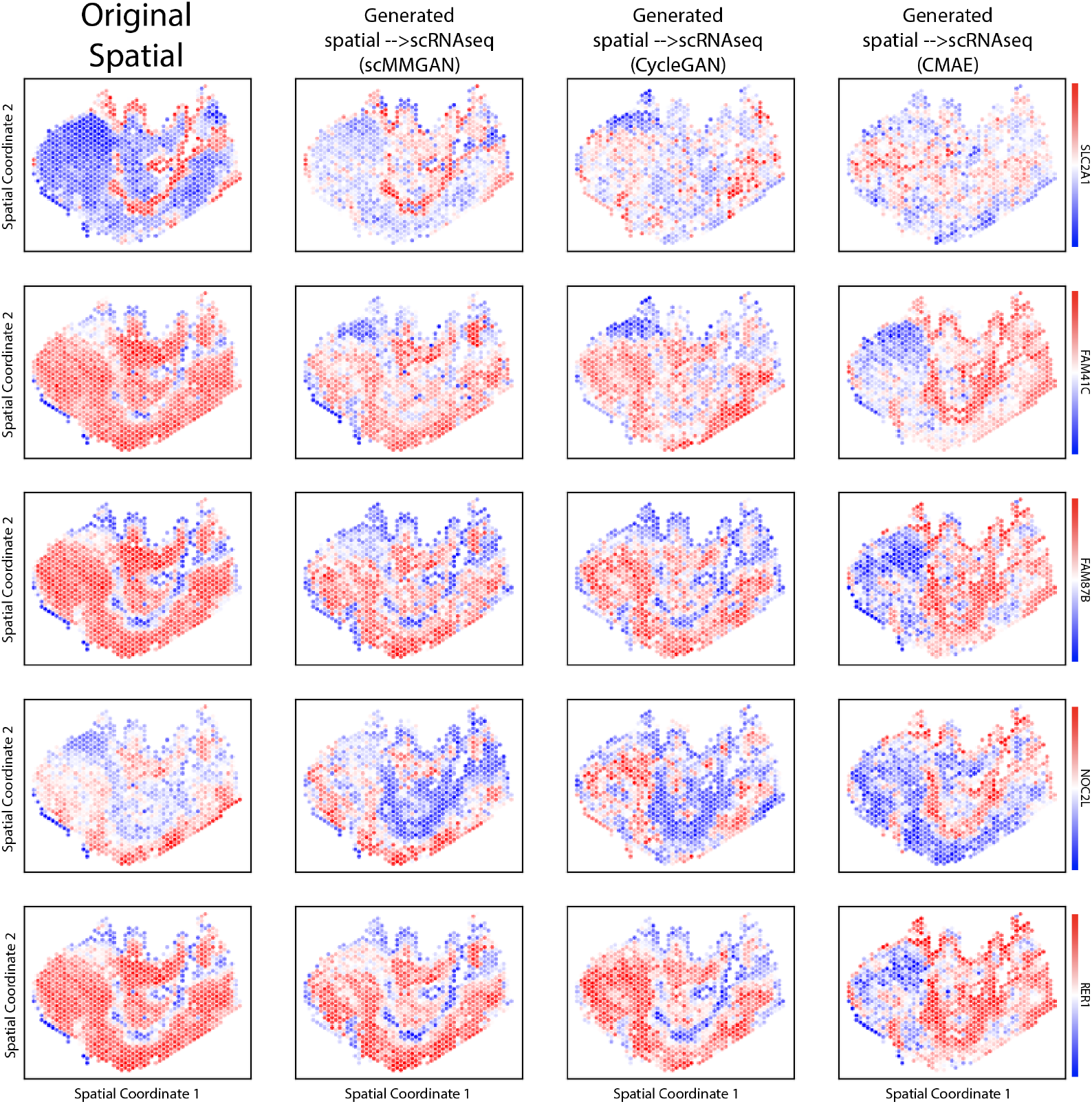
The x-axis and y-axis plotted are the raw measured spatial coordinates from the spatial RNAseq. The color is expression value, where we compare the original spatial RNAseq of a gene with each generated scRNAseq value of that gene for each method, showing scMMGAN best aligns the original and generated values.

In the first column of Figure 6, we plot the value of five genes across the original spatial coordinates in the spatial RNAseq data. Then, we plot the generated scRNAseq value of that gene for each spatial coordinate for each model in the subsequent columns, starting with scMMGAN. With FAM87b in the first row, we see scMMGAN’s generated values largely match the original spatial pattern, with some minimal changes that were necessary to match the target distribution as well. The CyleGAN matches much of the bottom half of the spatial coordinates, but in the top half maps some coordinates that were low in the gene to scRNAseq profiles that are high in the gene and vice versa. The CMAE has even less correspondence between the original spatial RNAseq value of the gene and the generated scRNAseq value.

The preservation of signals by scMMGAN and not the other methods has important consequences for downstream analysis. In the first row of Figure 6, we show the values of SLC2A1. This gene encodes the glucose transporter Type 1 (GLUT1) protein that is commonly upregulated in triple-negative breast cancers and is associated with high grade tumors, having been previously shown to be a potential driver of metastasis in a broad array of breast and other cancers [27, 28, 29, 30, 31]. Notably, in the spatial data, SLC2A1 activity has a strong spatial pattern in which areas in the tissue express it highly. With the scMMGAN mapping, the spatial observations with high SLC2A1 also have high expression in the generated scRNAseq data. With the other models, however, the SLC2A1-high spatial observations are mapped to SLC2A1-low scRNAseq cells. This important signal has been lost, and the downstream analysis that seeks to understand the differential spatial distribution and function will have lost this key gene signal. The scMMGAN mapping produces aligned data that best preserves the original signal.

The bottom row showing RER1 demonstrates another canonical situation motivating scMMGAN’s correspondence loss. This gene is roughly bimodally distributed with equal numbers of observations high and low in it. Because flipping two populations is often as easy as introducing a single negative sign into a single weight in a neural network layer, CMAE maps all spatial coordinates high in the gene to scRNAseq profiles low in the gene and vice versa. Only with scMMGAN’s correspondence loss is one of these equally-easy-to-learn mappings specified as preferable, with the training objective significantly lower for the one that doesn’t flip the populations as opposed to it being equal. This is further corroborated by the results of the quantitative experiments shown in Table 3.

**Table 3:**
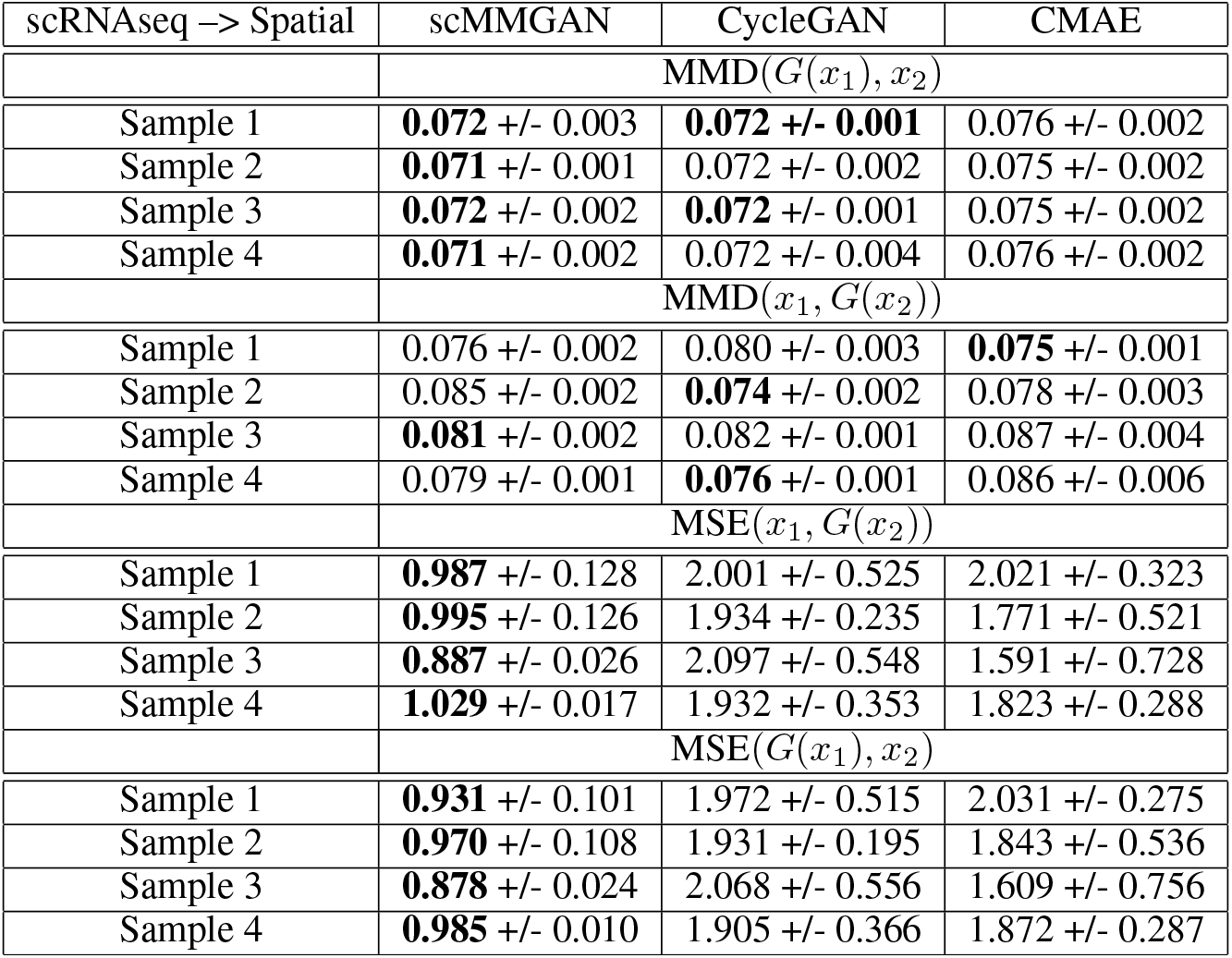
Quantitative measurement of how well the generated distributions match the target distribution (MMD) and how well they preserve correspondence with the original input distribution (MSE). While all methods match the target distribution reasonably (top 2 sections), only scMMGAN minimally alters the points in the alignment (bottom 2 sections). Statistics reported on both mapping directions and across 5 independent trials. The best or tied-for-best values are bolded.

#### Gene correlations

scMMGAN highlights the differences between the measurements of the two modalities by investigating the genes most highly correlated to a particular gene of interest in this system in both the original data versus the generated data. Specifically, as the spatial data is an aggregate measurement of multiple cells in the same proximate area in the tissue (not single cell), we can highlight some possibly spurious correlations by mapping them to the scRNAseq space and re-calculating the correlations.

Consider for example the glucose-transporter gene SLC2A1 that we have studied previously. If we look at the *n* most correlated genes in the original spatial data, any of them that have low correlation in the generated scRNAseq data are candidates for spurious data artifacts. Similarly, any of the *n* most correlated genes in the generated scRNAseq data that have low correlation in the original scRNAseq are candidates for novel associations found by the scMMGAN.

Choosing *n* = 700 and defining low correlation in the other space as being less than 0.1, we get a list of six genes that have spurious correlations: FRAT2, CAMK2A, CANX, LRRC66, ZMIZ1-AS1, MTRNR2L8. Then we have the following five genes that have been discovered by scMMGAN: AC092115.3, P2RX7, CCDC93, UTP25, BBS10.

Among these genes whose correlation to the glucose-transmitter SLC2A1 is discovered by scMMGAN, we see P2RX7, which has been identified as a precursor to glucose transporters in the literature [32]. This provides corroborating support in favor of the scMMGAN-discovered gene correlations.

## 4 Discussion

In this work we demonstrated that scMMGAN can align data from related experiments but different modalities in a way that best preserves the properties of the original cells through the learned mapping. The addition of the correspondence loss in scMMGAN’s architecture resolves the ambiguity created by only stating a distribution-level loss in learning a mapping. This holds across a wide array of data types and modalities, distribution shapes, and other settings that arise in practical biological experiments.

We have shown how scMMGAN can be used to measure uncertainty in the mapping and use injected stochasticity to gauge which information is unique to one of the modalities and which information is common between them. This can be used to not only answer questions about the cellular samples in an experiment, but also can be used to answer questions about the technologies and modalities themselves, in terms of their strengths and weaknesses.

Furthermore, we have shown how scMMGAN can be used to identify spurious correlations found in one modality as artifactual results as opposed to real findings. Similarly, we demonstrated scMMGAN’s ability to identify novel correlations that are not visible in an individual modality but become apparent when the data is mapped to another modality. In these ways, scMMGAN can be added to traditional analysis pipelines to uncover further insights from complicated, multi-modal experimental data.

### Limitations

There are limitations to the proposed approach that bear mentioning. Although GANs have been useful in mapping distributions, they suffer from key drawbacks. First, they are difficult to train because of the adversarial losses, which can lead to instability [33]. This instability means that the model can deteriorate from effective to ineffective quickly across training iterations. Second, they often suffer from mode collapse since they are not penalized by distribution-level losses to match the entire distribution [34]. The additional correspondence loss does not worsen these issues. We informally observe that early stopping, as a regularization, as well as our geometric loss helps mitigate these effects. But these effects may still be present in some contexts. Additionally, with our framework based on pairwise generators in each mapping direction, the number of generators necessary grows quadratically. This means that for a large number of input modalities to align, the networks would have to be made small or they would have to be trained separately. Finally, our geometry-based loss is not completely plug-and-play in the sense that we still require a choice of distance between data points. In cellular data, we used Euclidean distance to compute the manifold. However in other contexts, such as two image types, more complicated distances such as SSIM may be used [35, 36].

For this reason, we encourage continued evaluation of aligned results through external verification measures. For example, in this work we verified that known signals across genes and across cells are still preserved in the aligned data. Moreover, we point out that the novel gene correlations found by scMMGAN are potential discoveries that should be further investigated with experiments specifically designed for this.

## 5 Experimental Procedures

In this section we further expand on the model, experimental regimes, and implementations used in this work.

### 5.1 Code availability

An implementation of the scMMGAN model written in Python and Tensorflow which can be run on any user-loaded datasets is available at *github.com/KrishnaswamyLab/scMMGAN*.

### 5.2 Training Objectives

Here we elaborate on the training objectives used in the scMMGAN framework learning. We define the formulation considering a pair of domains, with the definitions extending to multiple domains accordingly. It is composed of distinct GAN networks, each with a generator network *G* with input *X* and output *X*′. We call each generator a *mapping* from the input domain to the output, or target, domain. Each generator attempts to make its output G(X) indistinguishable by *D* from *X*′. Denote the two datasets *X*_1_ and *X*_2_. Let the generator mapping from *X*_1_ to *X*_2_ be *G*_12_ and the generator mapping from *X*_2_ to *X*_1_ be *G*_21_. The discriminator that tries to separate true samples from *X*_1_ from the generated output of *G*_21_(*X*_2_) is *D*_1_ and the discriminator that tries to separate true samples from *X*_2_ from generated samples from *G*_12_(*X*_1_) is *D*_2_.

The loss for *G*_1_ on minibatches *x*_1_ and *x*_2_ is:

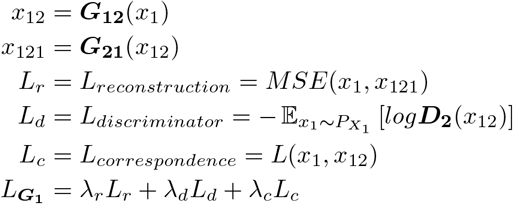

where MSE is the mean-squared error and L is the correspondence loss discussed previously. The hyperparameters λ_*r*_, λ_*d*_, and λ_*c*_ are chosen to balance the reconstruction, discriminator, and correspondence losses. These can be chosen by default to be λ_*r*_ = λ_*d*_ = λ_*c*_ = 1, but λ_*c*_ increased if the observed correspondences are low and λ_*r*_ increased if the observed reconstructions are not accurate.

Similarly, the loss for *G*_2_ is:

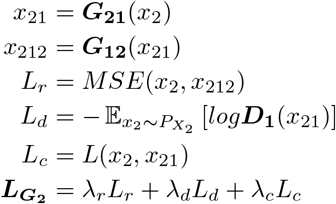

The losses for *D*_1_ and *D*_2_ are:

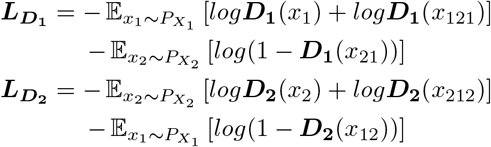

### 5.3 Calculation of correspondence loss

For the following notation, consider one of the datasets *x*_1_ and its representation after being mapped to the other domain, *G*(*x*_1_). First, matrixes of pairwise distances *D*_*x*_1__ and *D*_*G*(*x*_1_)_ are constructed.

Then, these are transformed into matrixes of pairwise affinities with an inverse-distance kernel *k*(*x_i_, x_j_*) = max(0,1 - ||*x_i_* - *x_j_*||_1_/*σ_i_*) where *σ_i_* is a k-nearest neighbor adaptive bandwidth. This kernel was necessary as the standard Gaussian kernel suffered from gradients that saturated and suppressed learning. With this kernel, we are able to perform diffusion geometry learning through gradient descent effectively.

These affinity matrixes are transformed into transition probability matrixes *P*_*x*_1__ and *P*_*G*(*x*_1_)_ through row-normalization. Powering these matrixes *t* times then represents taking *t* steps forward in the Markov chain. Let the eigenvectors of these two matrixes be *ϕ*_*x*_1__ and *ϕ*_*G*(*x*_1_)_, respectively. The *i^th^* row of *ϕ*_*x*_1__ and *ϕ*_*G*(*x*_1_)_ represents the diffusion coordinates of the point *x*_1*i*_ in the original space and *G*(*x*_1*i*_) in the generated space, respectively. Because we are only seeking to match low-frequency structure of the data, we use only the first *n_e_* eigenvectors of the data as an approximation. The eigenvectors are then rescaled to be between −1 and 1.

We also perform a check before comparing the eigenvectors of the original data *ϕ_x_* and those of the generated data *ϕ*_*G*(*x*)_. Because the direction of the eigenvectors can be switched, two datasets with equivalent intrinsic geometry could have eigenvectors that are either highly correlated or highly anticorrelated. To combat this, we calculate the correlation of each pair of eigenvectors before computing the loss and if the correlation is below a threshold *ε_r_* = –0.5 then we multiply the values of *ϕ*_*G*(*x*)_ by −1 before computing the loss. We then also compare the eigenvectors at different scales, by summing adjacent vectors and comparing the new combined representations that have half the number of vectors.

### 5.4 Noise-augmented Model

Here we detail the noise-augmented model used in the section about distinguishing unique and common information. The core idea is that by providing additional noise as input, the model will be able to use the stochasticity when necessary or ignore it if not. In other words, some generated values will have more certainty behind them in the model and others greater uncertainty. We experiment with this notion by introducing a slight modification of the scMMGAN framework: additional random input. We calculate the generator’s output as *G*(*x_i_*; *z*) where *z* ~ *N*(0,1) ∈ ℝ^*D*^, concatenating a draw from an isotropic normal distribution with the original input. The reconstruction and correspondence losses are then calculated as usual with just *x_i_*. This allows the model to create more stochasticity around regions of the space where there is not enough information to pin down precisely the correct alignment, while it can ignore the noise and create a deterministic mapping in regions of the space where there is enough information.

### 5.5 Invariance and Risk

In this section we connect our model in the domain alignment setting to existing literature on invariance and risk minimization [37]. Consider the domain alignment task as drawing a dataset 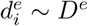 from a distribution *P*(*D^e^*) where the environment *ϵ* determines how observations manifest *d_i_* in *D_e_*. The domains *X*_1_ and *X*_2_ in our setting are drawn from *P*(*D^e^*), and then the datasets 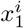, *i* = 1…*n*_1_ and 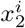, *i* = 1…*n*_2_ are drawn from the distributions *X*_1_ and *X*_2_. Thus, we have two sources of randomness from sampling to consider in learning the desired mappings *G*, one from the points sampled from the distribution and the other from the distribution sampled from the distribution over settings. We want to minimize the risk of alignment:

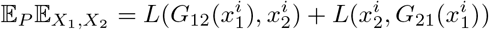

The inadequacy of just a cycle-consistency loss on its own should be obvious from this formation. While minimizing the cycle-consistency loss can optimize performance with respect to the expectation over *X*_1_ and *X*_2_, the formulation is equivalent for all datasets drawn from *P*. Thus, it is incapable of resolving correlations that are structurally related to the datasets versus those arising spuriously due to sampling. These ideas are key to the extension beyond just using cycle-consistency in the construction of domain mapping networks.

### 5.6 Diffusion Maps

Diffusion maps define a process of Markovian diffusion over a dataset, where a set of local affinities capture the intrinsic data geometry as quantified by diffusion distances [18]. They operate on a matrix of pairwise distances that is transformed into a matrix of pairwise affinities, here via the commonly used Gaussian kernel *k*(*x_i_, x_j_*) = exp{-||*x_i_* - *x_j_* ||^2^/*ϵ*}. A Markov chain transition matrix over the dataset *P* is constructed from the pairwise affinity matrix *A* via 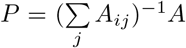. Powering the matrix *P^t^* represents taking *t* forward steps in the Markov chain. Diffusion maps are then defined as 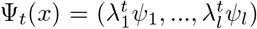 where λ_*i*_ and *ψ_i_* are the *i^th^* eigenvalue and eigenvector of *P* and *l* is a hyperparameter of the number of top eigenvectors to use. The diffusion map coordinates form a space where the Euclidean distance between points approximates the diffusion distance between those points.

In previous work, diffusion operators have been used in the context of multimodal data integration [38]. This has been done for the related tasks of visualizing and denoising, rather than mapping between, the datasets. The approach there differs from ours in that it relies on combining diffusion operators from different modalities through algebraic operations as opposed to our method which integrates them into a broader deep learning framework.

The diffusion maps are a key foundational notion used in the construction of the data geometry loss.

### 5.7 Geometry-preserving Correspondence Loss

We now further elaborate on a few points about the geometry-preserving correspondence loss introduced in this paper. We only enforce the correspondence loss on the first eigenvectors because this ensures that basic low-frequency signals are largely aligned while still allowing the flexibility of changing high-frequency signals that are more likely to be idiosyncratic to each domain. In practice we find using 10-20 eigenvectors work best.

We note that any changes to the data geometry would cause a mismatch here, and thus the ideal alignment would not drive this term in the objective all the way to zero unless the two datasets being aligned have identical geometry. Despite the goal not being zero correspondence loss, there are many different mappings that achieve comparably low GAN losses, and amongst them the ones with lower correspondence losses are preferable. This is why using it as a regularization to lightly guide the transformation in addition to the GAN loss can achieve the best performance overall.

### 5.8 Architecture and baselines

We compare scMMGAN to alternative baseline deep learning models used for alignment of this type: a CycleGAN, in order to motivate the need for the correspondence loss by showing the improper alignments obtained without it [9]; CMAE (Cross-Modal Autoenoder), an autoencoder-based model that uses separate encoders/decoders that learn to map into a shared space and then generates by crossing the encoder of one domain with the decoder of another [21]. These alternative methods use distribution-level losses to ensure the generated distribution matches the target distribution, but do not impose any loss on the representation of a point and its representation in the aligned domain. As a result, they can produce alignments which unnecessarily invert signals and change values of individual points.

With scMMGAN, we use a generator consisting of three internal layers of 128, 256, and 512 neurons with batch norm and leaky ReLU activations after each layer and a discriminator consisting of three internal layers with 1024, 512, and 256 neurons with the same batch norm and activations except with minibatching after the first layer [33, 39]. We use a correspondence loss coefficient of 10, cycle loss coefficient of 1, a learning rate of 0.0001, and a batch size of 256. As preprocessing steps prior to running each model on this dataset, we correct for dropout with the manifold smoothing method MAGIC [40], zero-center and unit scale each dimension, and reduce to fifty principle components. We use these architectures and hyperparameters in all subsequent experiments except where otherwise stated.

